# Single-cell and single-variant resolution analysis of clonal evolution in human liver cancer

**DOI:** 10.1101/2020.12.30.424907

**Authors:** Xianbin Su, Linan Zhao, Yi Shi, Rui Zhang, Qi Long, Shihao Bai, Qing Luo, Yingxin Lin, Xin Zou, Shila Ghazanfar, Kun Tao, Guoliang Yang, Lan Wang, Kun-Yan He, Xiaofang Cui, Jian He, Jiao-Xiang Wu, Bo Han, Na Wang, Xiaolin Li, Pengyi Yang, Shangwei Hou, Jielin Sun, Jean Y. H. Yang, Jinhong Chen, Ze-Guang Han

**Author notes:** Equal contributors (Xianbin Su, Linan Zhao, Yi Shi).

## Abstract

Genetic heterogeneity of tumor is closely related to clonal evolution, phenotypic diversity and treatment resistance. Such heterogeneity has been characterized in liver cancer at single-cell sub-chromosomal scale, and a more precise single-variant resolution analysis is lacking. Here we employed a strategy to analyze both the single-cell genomic mutations and transcriptomic changes in 5 patients with liver cancer. Target sequencing was done for a total of 480 single cells in a patient-specific manner. DNA copy number status of point mutations was obtained from single-cell mutational profiling. The clonal structures of liver cancers were then uncovered at single-variant resolution, and mutation combinations in single cells enabled reconstruction of their evolutionary history. A common origin but independent evolutionary fate was revealed for primary liver tumor and intrahepatic metastatic portal vein tumor thrombus. The mutational signature suggested early evolutionary process may be related to specific etiology like aristolochic acids. By parallel single-cell RNA-Seq, the transcriptomic phenotype was found to be related with genetic heterogeneity in liver cancer. We reconstructed the single-cell and single-variant resolution clonal evolutionary history of liver cancer, and dissection of both genetic and phenotypic heterogeneity provides knowledge for mechanistic understanding of liver cancer initiation and progression.

## Introduction

Tumor heterogeneity is closely related to treatment response and drug resistance [1], and such heterogeneity in hepatocellular carcinoma (HCC), the third leading cause of worldwide cancer-related mortality, has been extensively studied [2, 3]. Single-cell sequencing has recently evolved into a powerful tool to interrogate tumor heterogeneity [4, 5]. The gene expression pattern of HCC cells was revealed by comparison with a normal human liver cell atlas [6], and the heterogeneity of cancer stem cell population was also assessed [7, 8]. The tumor immune microenvironment is believed to be potential therapeutic target as revealed by several studies of the landscape and dynamics of T cells and other immune cells in HCC [9, 10]. It was found that higher microenvironment diversity is associated with worse patient survival [11], and therefore immune-phenotypical classification will facilitate therapy decision [12]. Recent investigation also showed close correlation of intra-tumor heterogeneity and adaptive immune response during HCC evolution [13].

Genetic variations are believed to be the basis of cancer initiation, and the genetic heterogeneity of HCC has been comprehensively studied in bulk samples [14]. The genetic heterogeneity of multiple lesions varied among HCC patients [15], and multi-region sequencing further demonstrated the intra-tumor spatial heterogeneity [16] and evolutionary trajectories of metastases in HCC [17]. A multi-region genomics and epigenomics analysis revealed the temporal heterogeneity of HCC by studying primary tumor and their paired relapses [18]. Recently, single-cell genomics is demonstrated to be useful for revealing the clonal structure and evolutionary feature of tumors [19–22]. It has been shown that 25 single cells from one HCC patient clustered into two sub-populations based on copy number variation (CNV) [23], and distinct evolutionary models were also proposed based on the CNV profiles of 96 single cells from three HCC patients [24].

While previous studies have provided much insight into genetic heterogeneity of HCC, a more precise single-cell dissection of clonal heterogeneity in HCC at single-variant resolution, such as single-nucleotide variation (SNV) or small insertion or deletion (INDEL), is still lacking. A recent micro-dissection analysis of 482 sites revealed the clonal dynamics in cirrhotic human livers [25], and whole genome sequencing of 48 hepatocytes from 12 human liver donors identified an age-related accumulation of somatic mutations [26]. As single-cell analysis of mutations in normal and cirrhotic liver has already been done, single-cell analysis of somatic mutations in liver cancer is thus needed for better understanding of its initiation and progression.

To address this, here we employed a single-cell strategy to analyze both the genomic mutations and transcriptomic changes in human HCC. Single-variant resolution clonal structures of HCC were constructed based on mutational profiles of single cells, and a common origin but independent evolutionary fates were found for primary and metastatic liver tumors. The clonal reconstruction of HCC at single-variant resolution can be directly related to functional alterations of specific genes, providing knowledge for mechanistic understanding. By analyzing high-resolution transcriptomes of single cells, we also observed relationship between genetic and transcriptomic heterogeneity, facilitating better understanding of genotype and phenotype linkage in human liver cancer.

## Methods

### Clinical specimens

The study was approved by the Ethnical Review Board of Shanghai Center for Systems Biomedicine, Shanghai Jiao Tong University. Both tumor and paratumor tissues were collected from a total of 5 HCC patients (HCC1, HCC2, HCC5, HCC8, and HCC9), including one (HCC8) with both primary tumor and the intrahepatic metastatic portal vein tumor thrombus (PVTT). The tissues were minced and digested in 0.05% collagenase IV (Sigma-Aldrich, C5138) at 37°C for 30 min with agitation. Centrifugations at 50× *g* for 2 min were repeated three times to enrich tumor cells, and a 40 μm-cell strainer (Corning, 352340) was used to remove undigested tissues. The single-cell suspensions were used for both single-cell whole genome amplification (WGA) and single-cell RNA-Seq (scRNA-Seq).

### Single-cell WGA and exome sequencing

Single-cell WGA was done on 10~17 μm or 17~25 μm C1 DNA-Seq IFC (Fluidigm, 100-5763 or 100-5764) via multiple displacement amplification method using illustra GenomiPhi V2 DNA Amplification Kit (GE Healthcare, 25660031). The WGA products from single cells were mixed according to their tissue origin, and the exonic regions were captured with Agilent SureSelect Human All Exon v7 Kit (Agilent, 5191-4005). Whole exome sequencing (WES) was done on illumina HiSeq platform with 2 × 150 bp mode. Variant calling, functional annotation, SNPs filtering, and mutational signature analysis were done as previously described [27].

### Selection of mutations for single-cell target sequencing

Putative clonal and subclonal mutations were selected from WES-derived mutation list (Additional file 1: Table S1), and the target sites were further narrowed down by checking mutation prevalence in cBioPortal collected HCC samples [28] and Gene Set Enrichment Analysis. A total of 57, 56, 57 and 64 mutation sites were selected for HCC1, HCC2, HCC9 and HCC8, respectively (Additional file 2: Table S2). HCC5 was not used as many mutations had low variant allele frequency (VAF) values, suggesting low tumor cell purity which was confirmed by scRNA-Seq. Primers were designed to amplify these sites from single-cell WGA product, and the amplification specificities of the primers were confirmed to ensure one PCR product for each primer pair.

### Single-cell target sequencing

A total of 480 cells were used for single-cell target sequencing, with 96, 96, 96 and 192 single cells from HCC1, HCC2, HCC9 and HCC8, respectively. WGA product from each single cell was used as PCR template for 5 parallel PCR reactions with ~10 primer pairs in each reaction. PCR cycling program was adopted from User Guide for Access Array System (Fluidigm, 100-3770) for even amplification of multiple targets.

For each single cell, PCR products from the 5 reactions were mixed for library preparation using FastStart High Fidelity PCR System dNTPack (Roche, 3553400001) and Nextera XT Index Kit (illumina, FC-131-2001). Single-cell libraries were mixed for sequencing on illumina NextSeq500 platform with 1 × 151 bp mode, and de-multiplexed based on their index combinations. The sequencing reads were mapped to GRCh37/hg19 with Tophat using default parameters, and numbers of the reads with reference or mutation site were counted.

### Single-cell mutational status and clonal structure analysis

The following criteria were used to determine the mutational status of a given target site: if reads <3, it was defined as site with no coverage; if reads >=3, more than 1 read with mutation, and VAF >=0.1, then it was defined as mutated site; other conditions were defined as reference site. All target sites in single cells were further confirmed by manual inspection in Integrative Genomics Viewer. We filtered genetic variations with poor coverage, and also removed those found in both paratumor and tumor cells which may be SNPs. Finally, 54/57, 55/56, 56/57 and 61/64 mutations were kept for HCC1, HCC2, HCC9 and HCC8, respectively. For cell filtering, we removed single cells with less than half of the sites covered, and 71/96, 60/96, 91/96, 177/192 cells were kept HCC1, HCC2, HCC9 and HCC8, respectively. We selected drop-out of normal allele for allelic drop-out (ADO) assessment.

The combination of mutations enabled inference of clones and clone-specific mutations in each patient, which could reconstruct their evolutionary relationship. For HCC9, 17 cells were found to be mixture of more than one cell based on VAF analysis and removed before clonal analysis. Besides single-cell clonal analysis based on the mutational status, we also used nucleotide sequences at each site for evolutionary phylogenetic tree reconstruction. The combined sequences from all target sites in the single cells were aligned, and maximum parsimony tree was constructed using MEGA-X.

### scRNA-Seq and data analysis

scRNA-Seq was done on 10~17 μm C1 mRNA-Seq HT IFC (Fluidigm, 101-4981). Single-cell libraries were pooled and sequenced on illumina HiSeq platform with 2 × 150 bp mode. The C1 mRNA Sequencing High Throughput Demultiplexer Script (Fluidigm) was used for de-multiplexing and generation of FASTQ files for each single cell. The sequencing reads were mapped to GRCh37/hg19 with Tophat using the default parameters, and the numbers of reads in each gene were counted. ERCC RNA Spike-in (Ambion, 4456740) was added as a technical control. SC3 was used for outlier identification and cell type identification [29]. Single-cell transcriptomes were filtered based on three parameters: total counts, total features, and ERCC count percentage. If all three parameters fall within 1 MAD (median of the absolute deviation) away from median then the cell was kept, and a total of 2064 cells from the 3200 cells passed the filtering. HCC8 samples were not analyzed because the size of tumor cell was out of the capture range of C1 HT IFC. Gene Ontology enrichment and transcriptional factor co-variance network analysis were done as previously described [27], and copy number inference from single-cell global transcriptional profiles was done with inferCNV software (https://github.com/broadinstitute/inferCNV).

## Results

### Overview of the single-cell analysis strategy of human liver cancer

A single-cell strategy was employed to analyze both the somatic mutations and transcriptomic profiles in 5 HCC patients (Fig. 1a,b and Additional file 3: Fig. S1). We mixed single-cell WGA products for WES, and 836 single cells from 6 tumor and 5 matched paratumor specimens were analyzed. Candidate clonal and subclonal mutations were selected for target sequencing in 480 single cells. High-resolution transcriptomes of 3200 single cells were generated from 4 tumor specimens, with high number of genes detected in single cells (Fig. 1c).

**Fig. 1.**
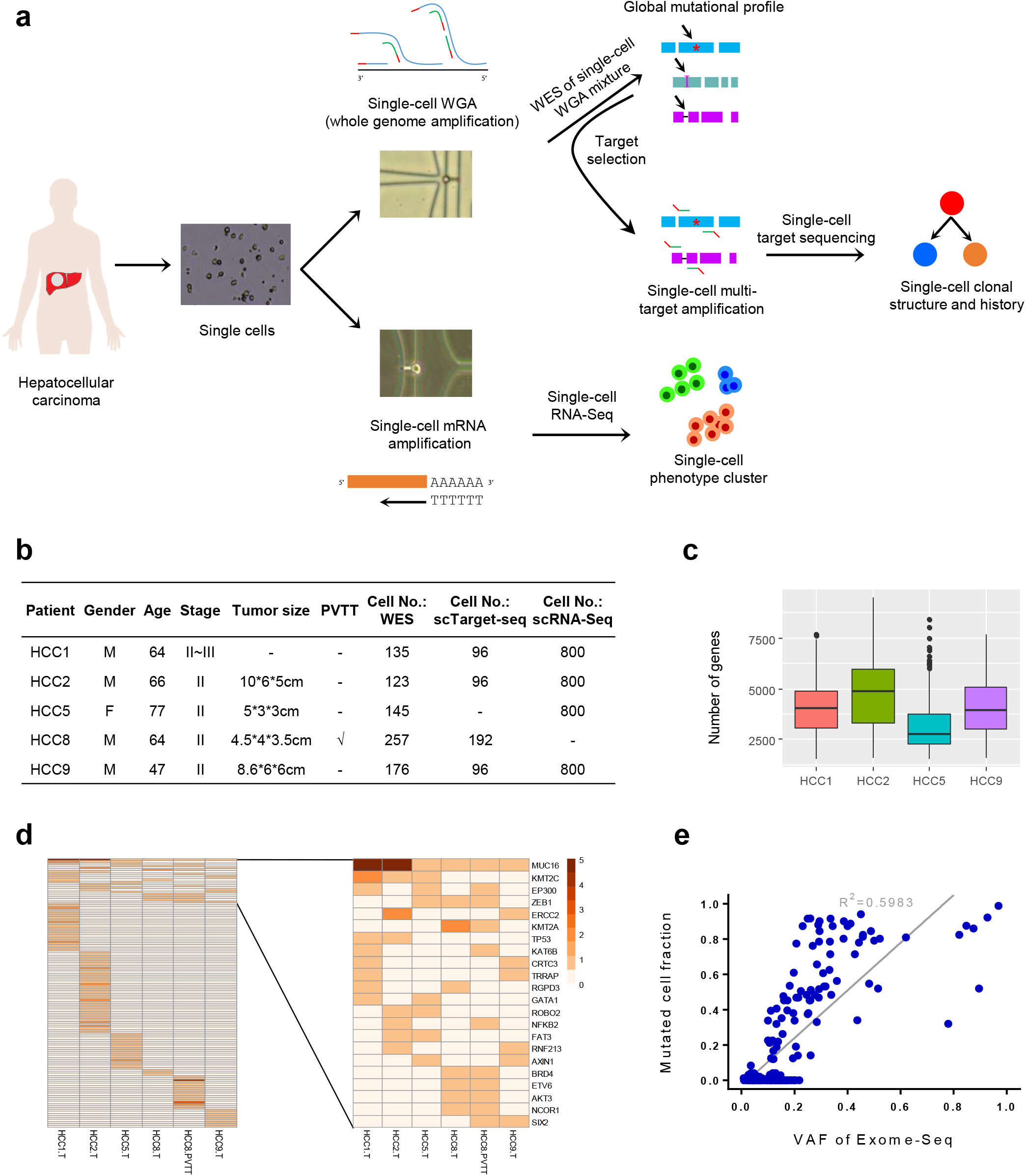
Overview of the single-cell analysis strategy of human liver cancer. **a** Experimental workflow. **b** Patient information and investigation summary. **c** Distribution of gene numbers detected in scRNA-Seq of each patient, with boxes showing median and 1^st^ and 3^rd^ quartiles, and whiskers showing 1.5 times of the inter-quartile range. **d** Heatmap showing the numbers of exonic mutations on driver genes by single-cell mixture WES. **e** Correlation plot between mutated cell fractions in single-cell target sequencing and WES-derived VAF values of mutations from HCC. Each dot represented a mutation, and the grey line represented linear regression with R^2^ shown.

Great inter-tumor genetic heterogeneity in human HCC were revealed by single-cell mixture WES. Around 200 ~ 900 exonic non-silent genetic variations were obtained for each tumor tissue (Additional file 1: Table S1, Additional file 3: Fig. S2a), and there were more tumor-private than shared mutations (Fig. 1d). Notably the paired primary tumor and PVTT of patient HCC8 only shared 143 mutations while acquired 123 and 354 private mutations, respectively, indicating occurrence of metastasis-private mutations during HCC progression (Additional file 3: Fig. S2b). For mutational signatures, the top nucleotide substitutions were C>T and T>C for patients HCC1, HCC2, HCC5 and HCC9, consistent with COSMIC mutational Signatures 12 and 30 (Additional file 3: Fig. S2c-e). The shared mutations in patient HCC8 primary tumor and PVTT showed prevalent T>A substitutions at CTG site, in agreement with COSMIC mutational Signature 22 (Additional file 3: Fig. S2d,e), which is related to carcinogens aristolochic acids [30]. Besides that, C>A substitutions reflecting COSMIC Signature 4 associated with tobacco mutagens were specifically higher in HCC8 primary tumor, implying independent accumulation of mutations in primary tumor and metastatic PVTT (Additional file 3: Fig. S2e).

Multi-target sequencing uncovered the genetic variations in HCC at single-cell resolution. Around 60 mutation sites were selected from each HCC patient for target sequencing (Additional file 2: Table S2), and high quality single-cell mutation data were obtained with ~8000× coverage for most covered sites. As expected, single cells from tumor tissues had mutations while those from paratumor tissues hardly had any mutation (Additional file 3: Fig. S3a). The fractions of cells mutated showed good correlations with WES-derived VAF values, indicating that the target sites were well detected (Fig. 1e). However, there were some mutations, mainly those with VAF values lower than 0.2, not detected in single-cell target sequencing, reflecting a difference between the pseudo-bulk and single-cell approaches. For the ADO issue, after filtering true homogenous mutations, except for HCC2, all other samples showed low drop-out rates (Additional file 3: Fig. S3b).

### Genomic DNA copy number changes of mutational sites in liver cancer

The single-cell target sequencing data allowed evaluation of genomic DNA copy number changes on the loci with point mutations. Most loci showed heterogeneous mutational status with VAF values fluctuating around 0.5, caused by uneven allelic amplification in single cells. However, there were homogenous mutations with VAF values centering on 1, such as *TP53, MUC16* and *FIP1L1* mutations in HCC1, and *AXIN1* mutation in HCC9 (Fig. 2a). Such homogenous mutations could be discriminated from the conditions where VAF values around 1 were caused by ADO in a fraction of cells, because most single cells harbored such mutations in the former condition. It was interesting that *CASC5* mutation in HCC9 and *NCOR1* mutation in PVTT of HCC8 both had VAF values around 0.75 in most single cells (Fig. 2a). This was unlikely caused by uneven allelic amplification at similar level in so many single cells, but should rather represent DNA copy number changes at these sites. Here these sites with VAF values of 0.75 could reflect possible amplification of the mutated allele.

**Fig. 2.**
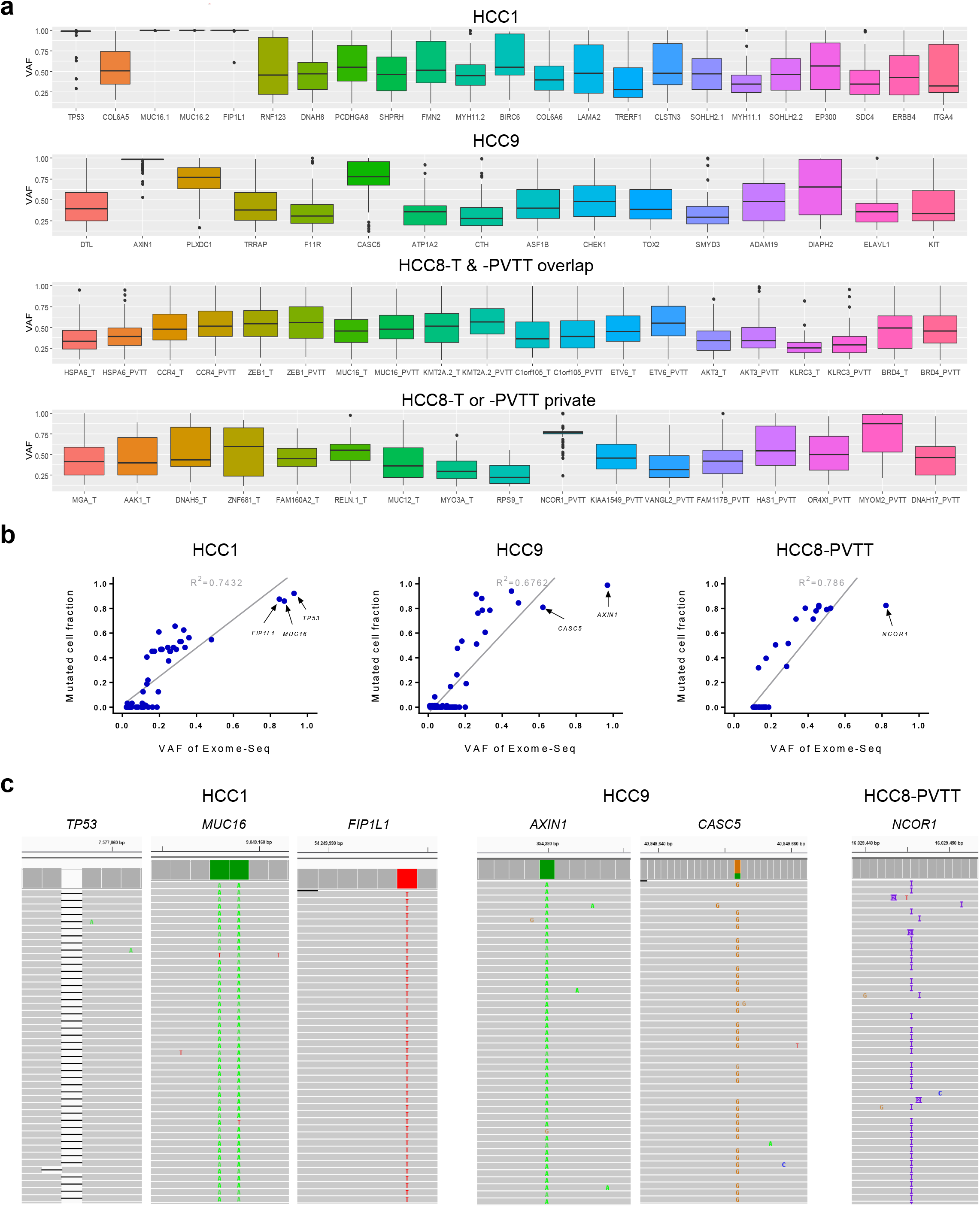
Copy number status of genomic loci with point mutations in liver cancer. **a** Distribution of VAF values of mutations in single cells, with boxes showing median and 1^st^ and 3^rd^ quartiles, and whiskers showing 1.5 times of the inter-quartile range. “_T” and “_PVTT” indicated data from primary tumor or PVTT. **b** Mutations with putative copy number changes highlighted in the correlation plot between mutated cell fractions and WES-derived VAF values. **c** Examples of sequencing reads with copy number changes at mutation sites.

Interestingly, the above-mentioned examples with copy number changes were all mutations on cancer-driving genes catalogued in the COSMIC Cancer Gene Census [31]. They were also observed in the correlation plot of mutated cell fractions and WES-derived VAF values (Fig. 2b), and further confirmed by inspection of the sequencing reads (Fig. 2c). The investigation of mutational copy number changes will be useful to distinguish tumor suppressors from cancer-promoting genes, and here *TP53* and *AXIN1* with homozygous mutations are both tumor suppressors.

### Single-cell clonal structures of liver cancer based on somatic mutations

The intra-tumor clonal heterogeneity of these HCC samples was then uncovered by mutational profiles from 399 high quality single cells after strict filtering. Both HCC1 and HCC2 exhibited a single-clone structure with limited heterogeneity, as most tumor cells had similar mutations (Fig. 3a,b). HCC1 and HCC2 both had functional clonal mutations in *TP53*, causing frame-shift after G294 in HCC1 and stop-gain at E198 in HCC2. By contrast, a multi-clone structure was observed in HCC9 (Fig. 3c,d), and also in primary tumor and PVTT from HCC8 (Fig. 4).

**Fig. 3.**
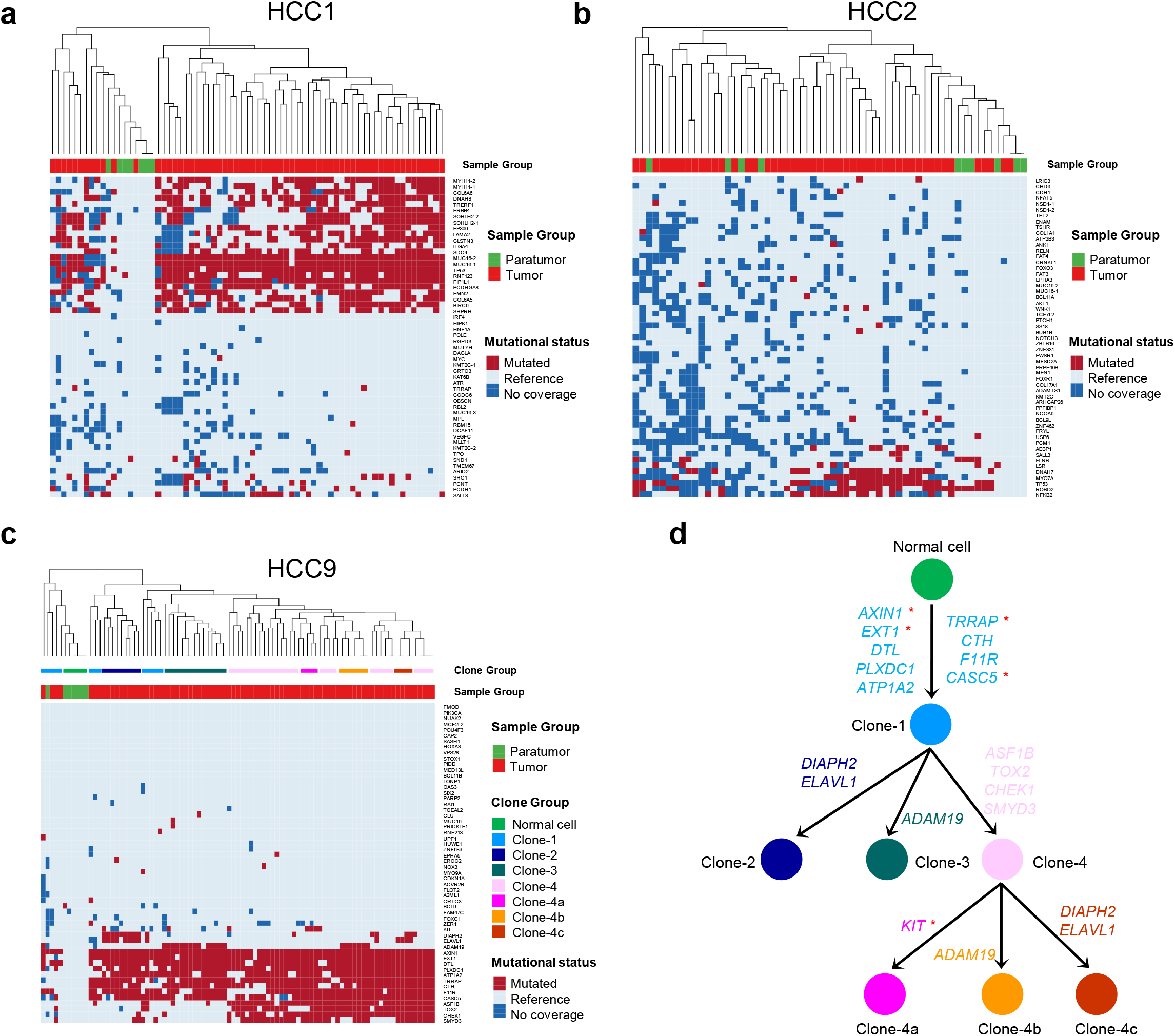
Single-cell clonal analysis of liver cancer based on point mutations. **a-c** Mutational status of SNV/INDEL sites in single cells from both paratumor and tumor tissues in HCC1 (**a**), HCC2 (**b**) and HCC9 (**c**). The inferred clone groups in HCC9 were also annotated. **d** Clonal evolution in HCC9 with genes mutated at each step shown. *: COSMIC Cancer Gene Census catalogued driver genes.

**Fig. 4.**
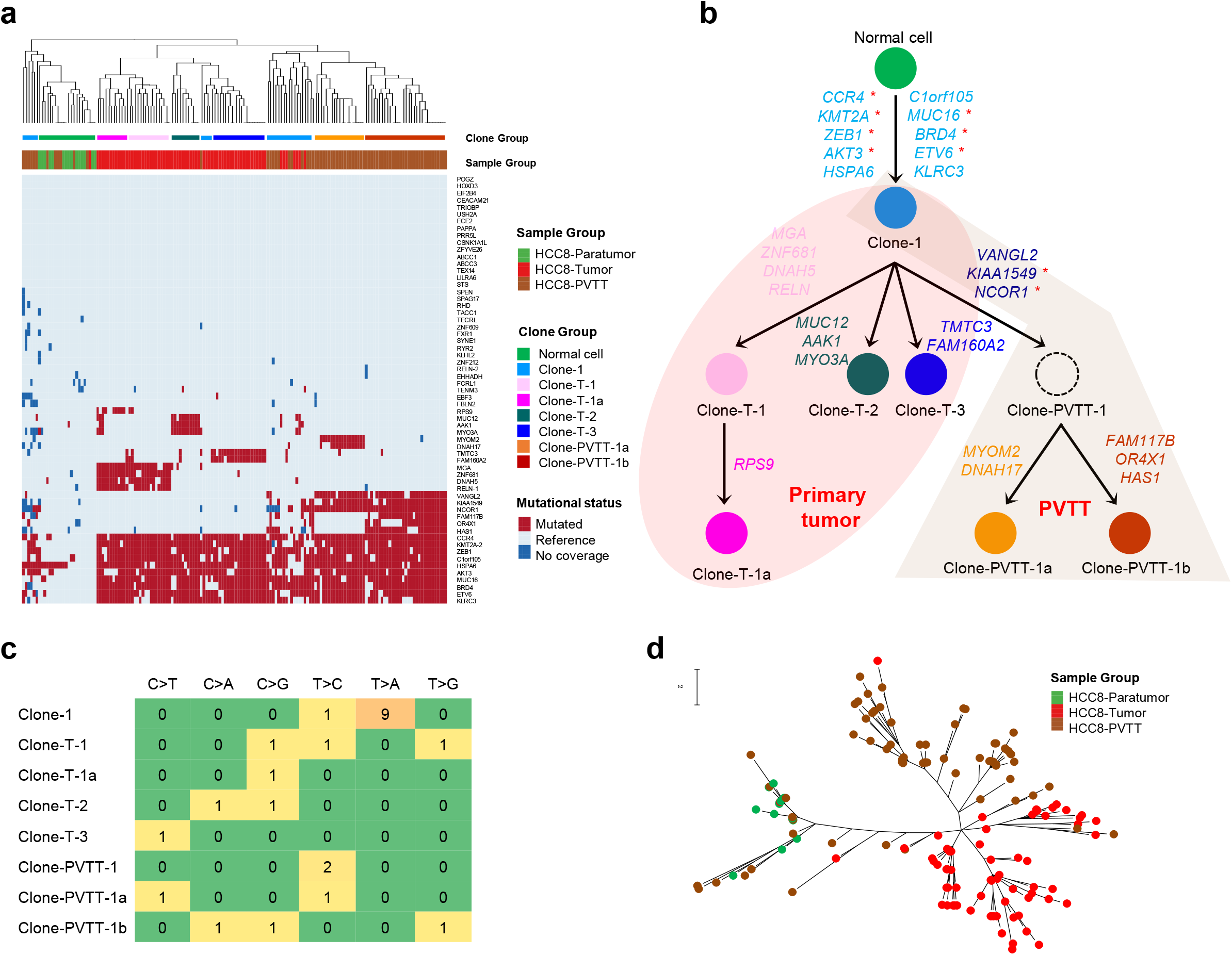
Common origin but independent evolution of primary and metastatic liver tumors. **a** Mutational status of SNV/INDEL sites in single cells from paratumor, primary tumor and PVTT tissues in HCC8. **b** Clonal evolution in HCC8 with genes mutated at each step shown. Dashed circle: virtual ancestor clone in PVTT. *: COSMIC Cancer Gene Census catalogued driver genes. **c** Statistics of nucleotide substitution types for clone-specific point mutations in HCC8 as shown in (**b**). **d** Maximum parsimony tree of single cells from HCC8 based on nucleotide sequences at the target sites. Scale bar: nucleotide substitution rate.

Compared with bulk sequencing-derived mutation list, the single-cell mutational profile clearly showed mutation combinations in each single cell. This enabled reconstruction of their evolutionary history despite that the single cells were collected at a fixed time point. In HCC9, the initiated cell was malignantly transformed to the founder Clone-1 as 9 genes (*AXIN1, EXT1, DTL, PLXDC1, ATP1A2, TRRAP, CTH, F11R* and *CASC5*) were mutated, where 4 of them are Census drivers: *AXIN1, EXT1, TRRAP*, and *CASC5* (Fig. 3d). The occurrence order of the 9 clonal mutations are not available based on the current data, however, and both successive accumulation and co-mutation might be possible approaches. Other clones were then derived from the founder Clone-1 by acquisition of extra subclonal mutations (Fig. 3d). For examples, the acquisition of *DIAPH2* and *ELAVL1* mutations led to Clone-2, the acquisition of *ADAM10* mutation led to Clone-3, and the acquisition of *ASF1B, TOX2, CHEK1* and *SMYD3* mutations led to Clone-4. Some of the cells in Clone-4 further evolved into 3 sub-clones by acquisition of more mutations. The results showed that clonal mutations may not necessarily be on driver genes, and interestingly majority of subclonal mutations were not on Census driver genes.

Single-cell mutational profiling clearly revealed the clonal status of given mutations in tumor cells. Based on their presence in all tumor cells, in some sub-clones, or in only a few cells or not detected, we can determine whether they are clonal, subclonal, or late-acquired rare mutations. While there are clear differences between mutated cell fraction values between clonal, subclonal and rare mutations, the WES-derived VAF values have overlaps for the three groups (Additional file 3: Fig. S4). This means that when a mutation is identified by bulk WES analysis, it might be difficult to determine its clonal status based on VAF values. This is reasonable as bulk sample VAF value is a reflection of combination of tumor purity that may be affected by cell constituents and VAF value in each tumor cell that may be affected by CNV as discussed above. The clonal status of mutations thus may only be reliably revealed by single-cell mutational profiling.

### Common origin but independent evolution of primary and metastatic liver tumors

To investigate the evolutionary relationship between primary and metastatic liver tumors, we carefully compared the paired primary tumor and PVTT from HCC8. Single cells from both primary tumor and PVTT shared mutations in 10 genes, where 7 of them are Census drivers: *CCR4, KMT2A, ZEB1, AKT3, MUC16, BRD4* and *ETV6* (Fig. 4a,b). Clone-1 with only the 10 shared mutations probably represent a common origin, and most other cells from both tumor tissues had divergent extra mutations, implying an early stage metastasis followed by independent evolution (Fig. 4b). Two clones within PVTT, Clone-PVTT-1a and Clone-PVTT-1b, shared PVTT-clonal mutations on *VANGL2, KIAA1549* and *NCOR1*, while also possessed other clone-private mutations. All PVTT-privately mutated genes are not Census drivers except PVTT-clonal *KIAA1549* and *NCOR1*, which have been suggested to be closely related to tumorigenesis [32, 33]. This suggested that mutations on the two genes might be metastasis-related early genetic events. The four clones within primary tumor, Clone-T-1, Clone-T-1a, Clone-T-2 and Clone-T-3, had other distinct mutations, and none of those mutated genes are Census drivers.

The clonal reconstruction at single-cell level could unveil nucleotide substitution pattern at each evolutionary step, which is relevant to HCC etiology and progression. Interestingly, nine out of ten mutations during the transformation of normal cell to the founder Clone-1 were T>A substitutions known to be related to carcinogen aristolochic acids, while the mutations from other clones had no common substitution pattern (Fig. 4c). This suggested that early genetic event during tumorigenesis may be related to specific etiology, while later evolutionary diversification may be complex. The finding was consistent with previous suggestion of early rather than late exposure to aristolochic acids for HCC development by bulk temporal mutational spectra analysis [34].

The phylogenetic tree based on target nucleotide sequences also showed a common origin of the single cells from primary tumor and PVTT, and the branches of single cells from both tumor tissues were consistent with the observed multi-clone structure in HCC8 (Fig. 4d). The proposed early metastasis in HCC by single-cell analysis is consistent with recent observation of early metastatic seeding in other tumor types [35]. As metastasis is an evolutionary process closely related to therapeutic vulnerabilities and drug resistance, the common origin but independent evolutionary fates for primary tumor and PVTT revealed at single-cell resolution in HCC might facilitate future treatment design.

### Genetic heterogeneity related to single-cell transcriptomic phenotypes

As the link of genotype and phenotype is useful for better understanding of tumor [36], here we conducted parallel scRNA-Seq analysis of HCC, which assigned single cells into macrophages, B cells, T cells and tumor cells (Fig. 5a and Additional file 3: Fig. S5). The inter-tumor and intra-tumor genetic heterogeneity in HCC were generally consistent with phenotypic heterogeneity. Tumor cells from 4 patients formed separate clusters, illustrating tumor-specific expression of genes with distinct Gene Ontology enriched terms and transcriptional factor networks, as well as heterogeneous expression of stem/progenitor markers like *CD24, ANPEP, SOX9, GPC3, AFP, EPCAM, LGR5*, etc. (Fig. 5b,c and Additional file 3: Fig. S6a,b).

**Fig. 5.**
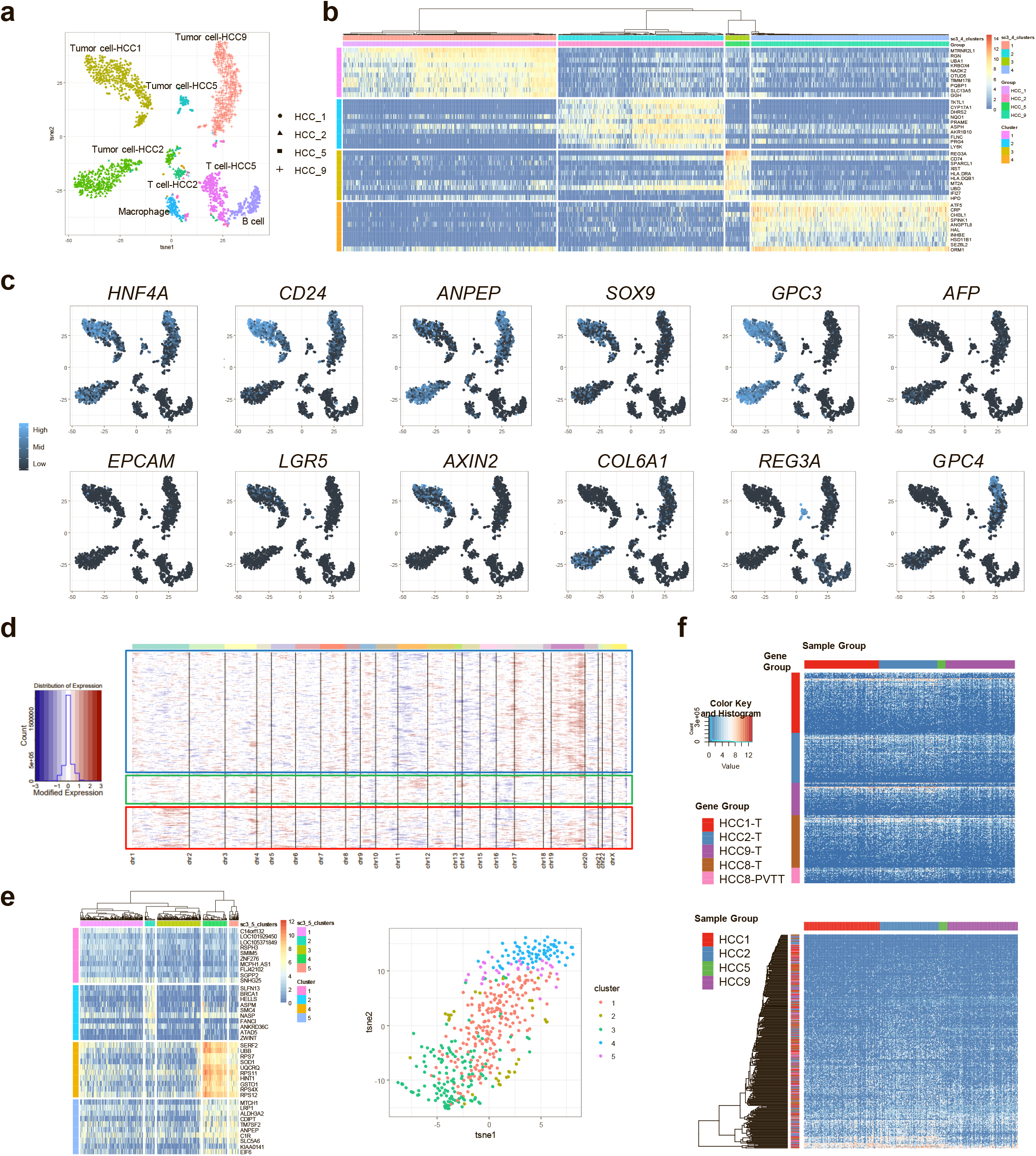
Link between genetic and phenotypic heterogeneity in liver cancer. **a** Cell clusters in scRNA-Seq analysis. **b** Heatmap showing patient-specific tumor marker genes. **c** The expression patterns of representative markers. **d** Copy number changes inferred from global transcriptomic profile of single cells in HCC9, with boxes highlighting sub-populations. **e** Heatmap (*left*) and t-SNE plot (*right*) showing the sub-populations in HCC9 based on differentially expressed genes. **f** The expression patterns of genes mutated in each HCC sample in scRNA-Seq data. Genes were grouped by tissue origin in the upper panel, and clustered by expression patterns in the lower panel.

For intra-tumor heterogeneity, the sub-chromosomal scale copy number changes inferred from single-cell global transcriptomic profiles in HCC1 or HCC2 were quite similar in each patient, but transcriptomic heterogeneity could still be observed among 3 ~ 5 sub-populations in each tumor based on differentially expressed genes, consistent with the single-clone structure and limited genetic heterogeneity evidenced at single-variant resolution (Additional file 3: Fig. S6c,d). For HCC9, 3 or 5 sub-populations were identified in tumor cells by copy number inference from global transcriptomic profile or differentially expressed genes (Fig. 5d,e), and the relatively higher phenotypic heterogeneity in HCC9 was also consistent with its higher genetic heterogeneity.

To understand the link between genetic mutations and transcriptomic changes, we checked the expression profiles of tumor-specific mutations. Interestingly, the genes mutated specifically in each liver tumor showed similar expression patterns among single cells from different patients, and clustering of those mutated genes using expression pattern exhibited a mixture of patient origin (Fig. 5f). This suggested that the tumor-specific mutations in HCC might cause phenotypic heterogeneity by altering the expressions of other genes rather than their own. The direct link between genetic and phenotypic heterogeneity of HCC, however, still await further clarification by future techniques that could detect point mutations and RNA transcripts simultaneously in a large number of single cells.

## Discussion

This study presented single-cell reconstruction of clonal evolution in liver cancer at single-variant resolution, and great inter-tumor and intra-tumor genetic heterogeneities were evidenced with single- or multi-clone structures observed in different patients. As SNV/INDELs are directly linked to functional alterations of specific proteins, a single-variant resolution tumor clonal structure will provide more precise information than CNV-based analysis [37]. Here the mutations used in single-cell clonal analysis were not randomly selected or from known gene panels, but from pseudo-bulk sequencing in a highly efficient tumor-specific manner. This will ensure the most relevant mutations were included for accurate clonal reconstruction in HCC.

Single-cell mutation combinations enabled reconstruction of the clonal evolutionary history, which is different from multi-region sequencing-derived tumor trees that may not represent genuine phylogenies [38]. Single-cell mutational analysis avoids the issue of tumor purity which may interfere with clonal inference in bulk analysis. The reliable clonal reconstruction at single-cell level could also unveil nucleotide substitution patterns at each step, providing cues for disease etiology [34]. Moreover, the single-cell approach tells not only the mutational status but also copy number changes for a given mutation site, facilitating full assessment of the combined functional effect.

It is interesting that most mutations on Census drivers are clonal mutations during the first step of normal cell transformation to founder tumor clone, or during early step of metastasis, as seen in HCC9 and HCC8. This indicated that the key genetic events occurred early during tumorigenesis, and at the time of tumor tissue collection the clonal structure only reflected most recent evolutionary scenario. To capture early genetic event, such as the mutational order of driver genes, we have to overcome the challenges of early diagnosis and temporal sampling in solid tumors in the future.

A limitation of this study is that single-cell target sequencing and scRNA-Seq are not done simultaneously in the same single cells, which prevents further linking genetic clones and phenotypic clusters. There has been integrated genetic and transcriptional analysis in leukemia by targeting somatic mutations on cDNA [39], but direct application to currently non-full-length scRNA-Seq may not be straightforward as the data are generally sparse and the profiled regions too short to harbor mutations. Analysis of the phenotypic heterogeneity based on scRNA-Seq may have the issue of accurate sub-population number identification [40], but here the relatively higher inter-tumor and lower intra-tumor phenotypic heterogeneity were found to be consistent with genetic level heterogeneity. The expression patterns of mutated genes also indicated that mutations were affecting other genes rather than themselves to cause phenotypic heterogeneity in HCC.

## Conclusion

In summary, here we reconstructed the clonal evolutionary history of HCC based on single-cell point mutations, and revealed a common origin but independent evolutionary fates for primary and metastatic liver tumors. The single-cell dissection of genetic and phenotypic heterogeneity provides knowledge to understand liver cancer initiation and progression.

## Abbreviations

ADO: allelic drop-out
CNV: copy number variation
HCC: hepatocellular carcinoma
INDEL: insertion or deletion
PVTT: portal vein tumor thrombus
scRNA-Seq: single-cell RNA-Seq
SNV: single-nucleotide variation
VAF: variant allele frequency
WES: whole exome sequencing
WGA: whole genome amplification.

## Acknowledgements

We would like to thank Prof. Gangcai Xie and Dr. Yanfeng Liu for proofreading the manuscript and providing valuable comments. This work is supported in part by National Natural Science Foundation of China (81802806), China National Science and Technology Major Project for Prevention and Treatment of Infectious Diseases (2017ZX10203207), Key Laboratory of Systems Biomedicine (Ministry of Education) Grant (KLSB2020QN-04), and Interdisciplinary Program of SJTU (ZH2018ZDA33).

## Data availability

The sequencing data have been deposited in NCBI GEO and SRA database under accession numbers GSE146115 and PRJNA606993.

---

**Supplementary Fig. 1.**
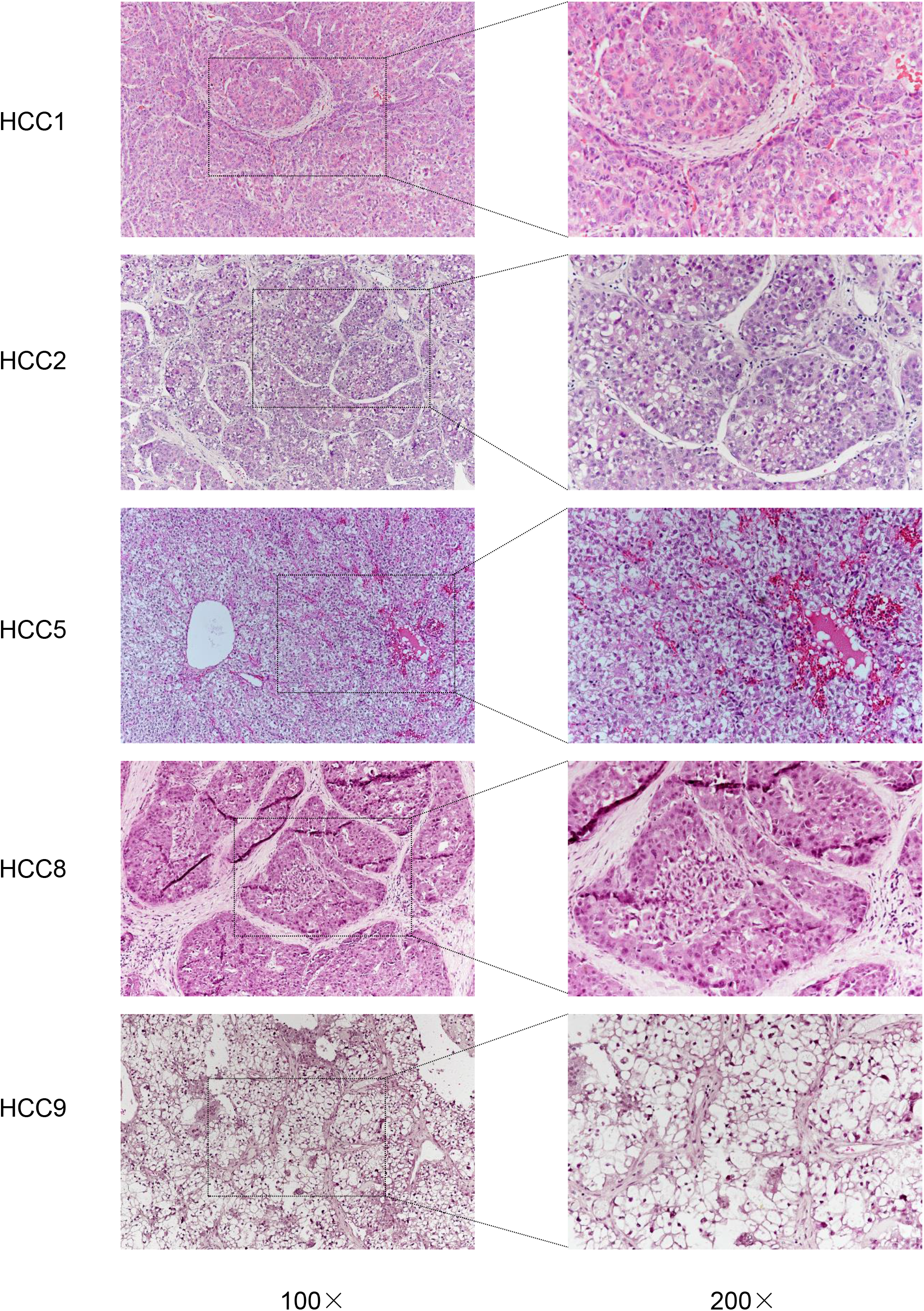
H&E staining of HCC tumor tissues.

**Supplementary Fig. 2.**
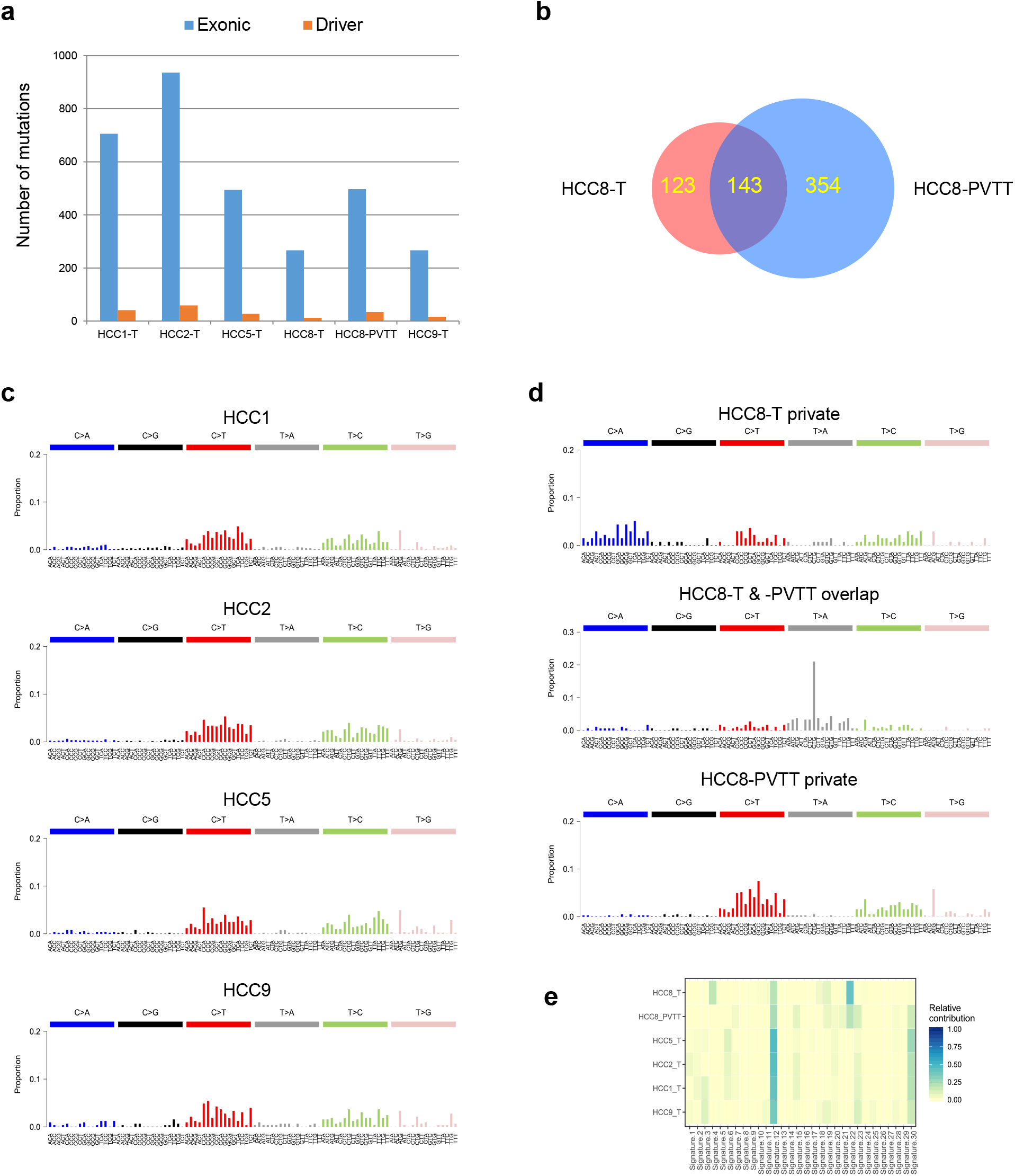
Single-cell mixture WES revealed inter-tumor genetic heterogeneity of HCC. **a** Number of exonic mutations and mutations on driver genes catalogued in the COSMIC Cancer Gene Census in HCC tumor tissues. **b** Overlap of exonic mutations in primary tumor and PVTT of HCC8. **c** Mutational spectra of genetic variations in tumor tissues from HCC1, HCC2, HCC5 and HCC9. **d** Mutational spectra of genetic variations shared or privately detected in primary tumor and PVTT of HCC8. **e** Contributions of COSMIC signatures to the HCC mutational spectra.

**Supplementary Fig. 3.**
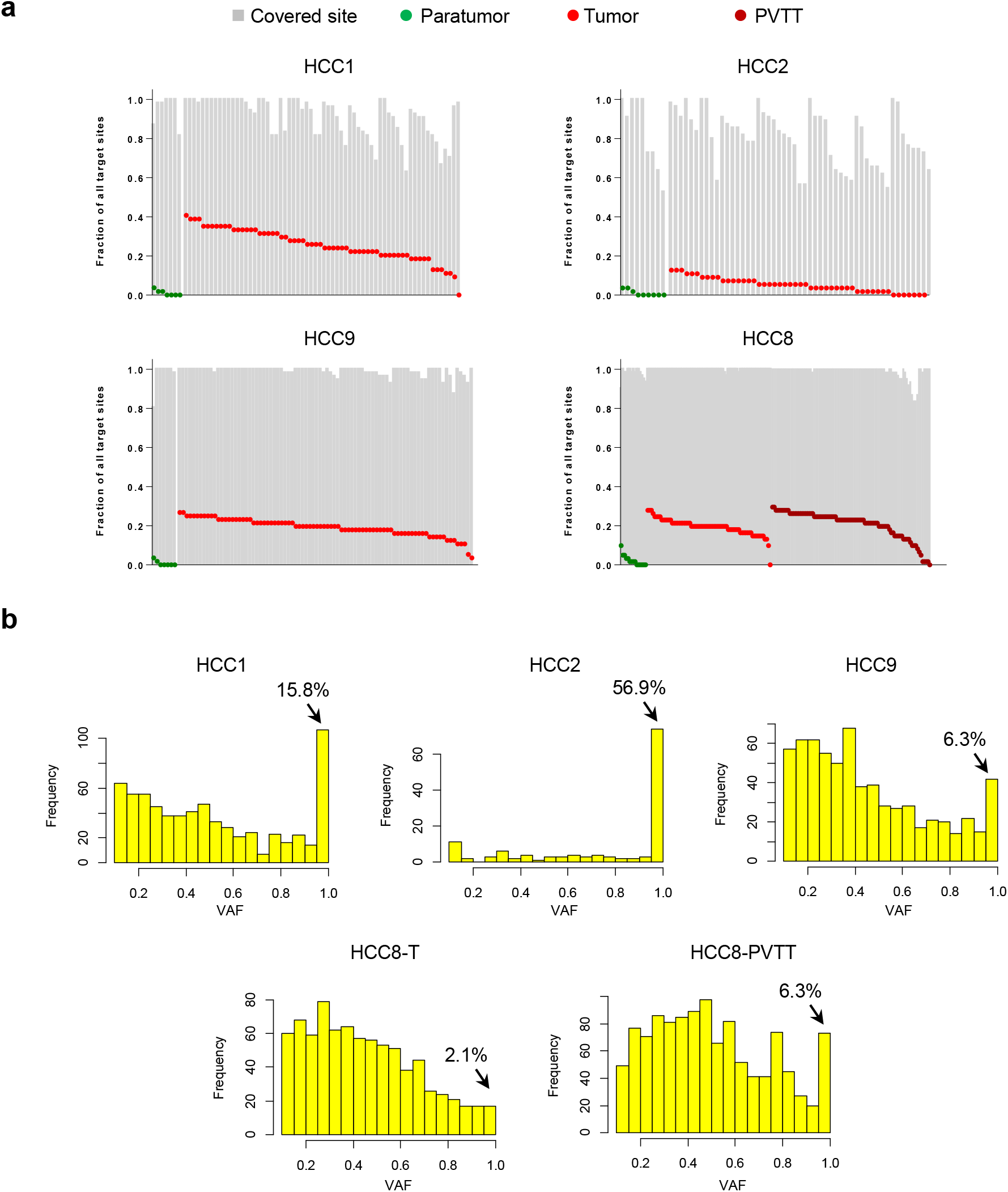
Quality assessment of single-cell target sequencing data. **a** Proportions of covered and mutated sites in each single cell in the 4 HCC patients. **b** Histograms showing the distributions of VAF values in single cells among each HCC sample, after filtering those values below 0.1. The percentage of variations with VAF values between 0.95~1 were shown in each sample for allelic drop-out assessment.

**Supplementary Fig. 4.**
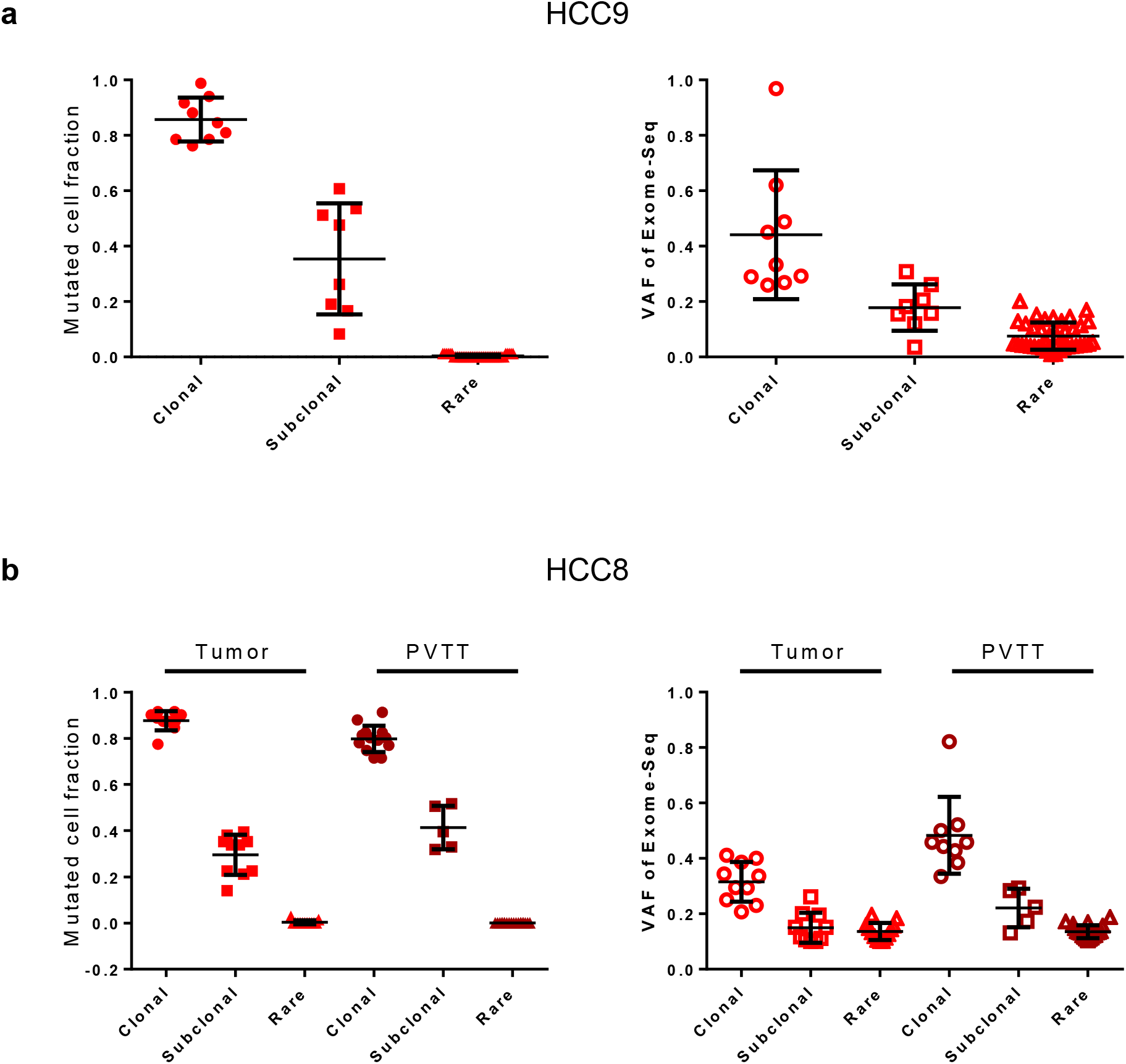
Comparison of mutated cell fraction and VAF values of clonal, subclonal and rare mutations in HCC9 (**a**) and HCC8 (**b**). Mutated cell fraction values are derived from single-cell mutational profiling, and VAF values are derived from single-cell mixture WES. The mutations were grouped into clonal, subclonal or rare mutations based on their clonal status among single cells. The mean and standard deviation are shown for each group. Both primary tumor and PVTT were included for HCC8.

**Supplementary Fig. 5.**
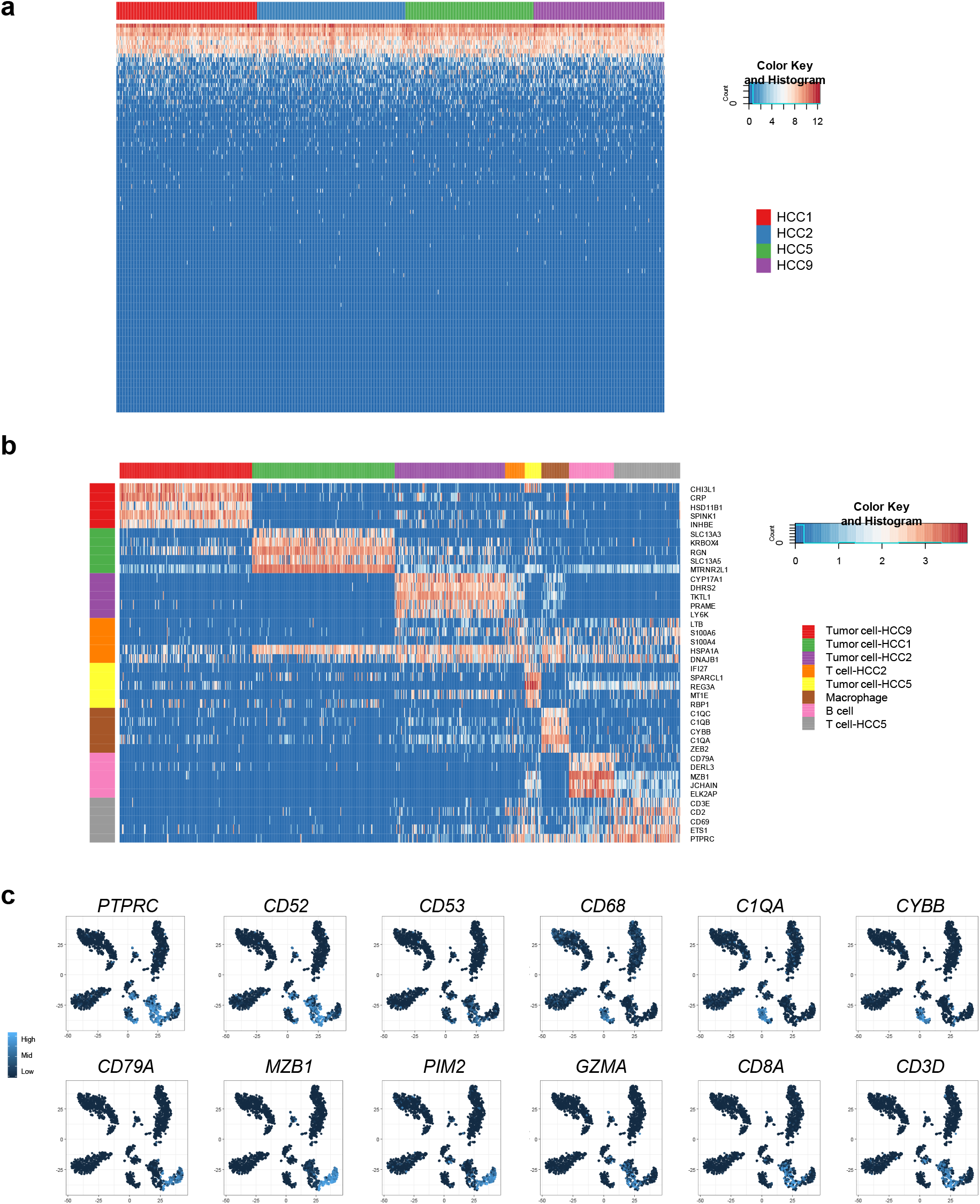
scRNA-Seq revealed the constituent cell types of HCC. **a** Heatmap showing the expression of ERCC RNA Spike-ins. Each row represented a spike-in, and each column represented a single cell. **b** Heatmap showing the genes specifically expressed in each cell cluster. **c** The expression pattern of representative immune markers.

**Supplementary Fig. 6.**
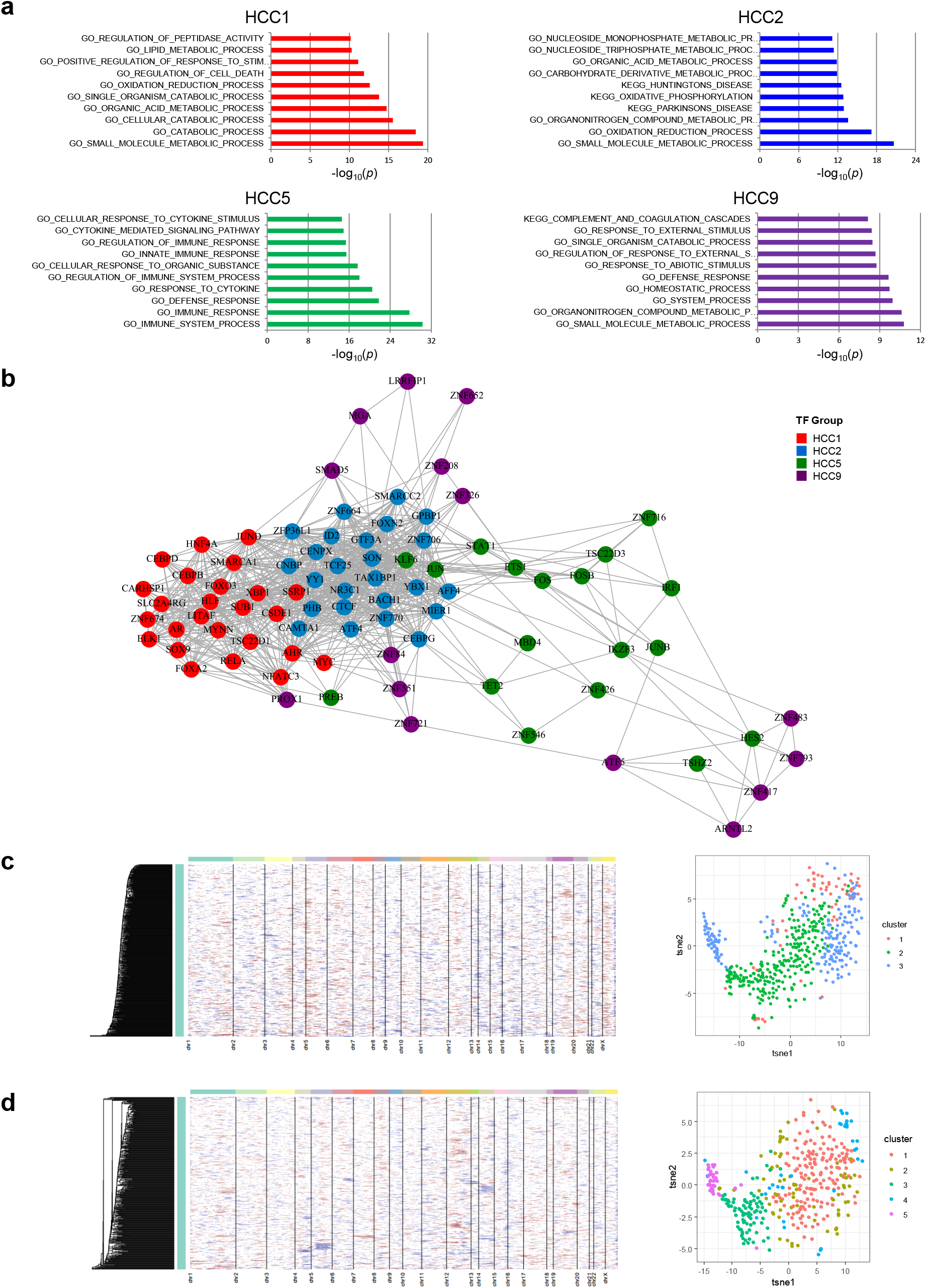
scRNA-Seq revealed the inter-tumor and intra-tumor heterogeneity of HCC. **a** The top 10 GO enrichment items for the genes specifically expressed in each tumor cluster. **b** Transcriptional factor covariance network of the HCC scRNA-Seq data. **c,d** The copy number changes inferred from global transcriptomic profiles of single cells (*left*) and the sub-populations based on differentially expressed genes (*right*) from HCC1 (**c**) and HCC2 (**d**).

